# A computationally tractable birth-death model that combines phylogenetic and epidemiological data

**DOI:** 10.1101/2020.10.21.349068

**Authors:** Alexander E. Zarebski, Louis du Plessis, Kris V. Parag, Oliver G. Pybus

## Abstract

Inferring the dynamics of pathogen transmission during an outbreak is an important problem in both infectious disease epidemiology. In mathematical epidemiology, estimates are often informed by time series of confirmed cases, while in phylodynamics genetic sequences of the pathogen, sampled through time, are the primary data source. Each data type provides different, and potentially complementary, insight; recent studies have recognised that combining data sources can improve estimates of the transmission rate and number of infected individuals. However, inference methods are typically highly specialised and field-specific and are either computationally prohibitive or require intensive simulation, limiting their real-time utility.

We present a novel birth-death phylogenetic model and derive a tractable analytic approximation of its likelihood, the computational complexity of which is linear in the size of the dataset. This approach combines epidemiological and phylodynamic data to produce estimates of key parameters of transmission dynamics and the number of unreported infections. Using simulated data we show (a) that the approximation agrees well with existing methods, (b) validate the claim of linear complexity and (c) explore robustness to model misspecification. This approximation facilitates inference on large datasets, which is increasingly important as large genomic sequence datasets become commonplace.

**Author summary:** Mathematical epidemiologists typically studies time series of cases, ie the *epidemic curve*, to understand the spread of pathogens. Genetic epidemiologists study similar problems but do so using measurements of the genetic sequence of the pathogen which also contain information about the transmission process. There have been many attempts to unite these approaches so that both data sources can be utilised. However, striking a suitable balance between model flexibility and fidelity, in a way that is computationally tractable, has proven challenging; there are several competing methods but for large datasets they are intractable. As sequencing of pathogen genomes becomes more common, and an increasing amount of epidemiological data is collected, this situation will only be exacerbated. To bridge the gap between the time series and genomic methods we developed an approximation scheme, called TimTam, which can accurately and efficiently estimate key features of an epidemic such as the prevalence of the infection and the effective reproduction number, ie how many people are currently infected and the degree to which the infection is spreading.

## Introduction

Estimating the prevalence of infection and transmission dynamics of an outbreak are central objectives of both infectious disease epidemiology and phylodynamics. In mathematical epidemiology, a time series of reported infections (known as the epidemic curve) is combined with epidemiological models to infer key parameters, such as the basic reproduction number, ℛ_0_, which is a fundamental descriptor of transmission potential [21, 53]. In phylodynamics, as applied to infectious disease epidemiology, phylogenies reconstructed from pathogen genetic sequences sampled over the course of an outbreak are used to estimate the size and/or growth rate of the infected population (eg [7, 30]).

Combining data from multiple sources has the potential to improve estimates of transmission rates and prevalence [9, 22, 33]. However doing so raises substantial technical challenges [23]. As a result phylogenetic and epidemiological inference methods have been developed and examined largely in isolation of each other [38, 46].

The two main frameworks for phylodynamic inference use the phylogenetic birth-death (BD) model, which estimates the rate of spread of the pathogen (eg [29, 39]), and the coalescent process, which estimates the effective size of the infected population (eg [26, 45]). Within the coalescent framework, a phylogeny reconstructed from sampled sequences is related to the effective size of the infected population and assumes that the fraction of the population that has been sampled is small [26]. This relationship, when interpreted under a suitable dynamical model, allows the inference of epidemic dynamics [16, 17]. Both deterministic and stochastic epidemic models have been fitted to sequence data [16, 18, 55], providing estimates of prevalence and ℛ_0_. [14] introduced an additional way to model effective population sizes, by considering the association between effective population size and time-varying covariates. [33] showed that combining sequence data with an epidemic time series could allow inference of not just the epidemic size but also its growth parameters. However, this approach treated the epidemic time series as being independent of the sequence data, an approximation which only holds when the number of sequences is small relative to the outbreak size. Previously, coalescent models have neglected the informativeness of sequence sampling times, although recent work has found estimates of the effective size can be improved substantially by incorporating sampling times (eg [27, 42]).

In the BD framework, births represent transmission events and deaths represent cessation of being infectious, eg due to death, isolation or recovery [50]. [48] extended this by modelling serially-sampled sequences as another type of death event. This approach was extended by [25], who linked the BD process to a stochastic epidemic (SIR) model under strong simplifying assumptions. The resulting model improved estimates of ℛ_0_ and provided the first means of inferring the number of unsampled members of the infected population (via estimates of epidemic prevalence). Deterministic SIR models have also been used in both BD [11] and coalescent frameworks [16].

[51] relaxed the assumptions in [25]’s model. This was made possible via the use of a particle-filter approach which enabled joint analysis of both sequence and epidemic time series data. While the particle-filter represents a comprehensive approach to fusing epidemiological and phylogenetic data, it is computationally intractable, relying on intensive simulation, which can limit its application. Data augmentation also provides a powerful approach to the inference problem, but again relies on intensive simulation [3].

Recently, [49] and [47] developed numerical schemes for computing the same likelihood, thereby facilitating equivalent estimation. Their methods have a smaller computational overhead, but still requires calculations that have a quadratic computational complexity, ie grow with the square of the size of the dataset. Moreover, the approximation used can be numerically unstable under certain conditions [1].

To the best of our knowledge, there is currently no existing phylogenetic inference method, in either the BD or coalescent frameworks, that can (i) formally combine both epidemiological and sequence data, (ii) estimate the prevalence of infection and growth rate, and (iii) be applied practically to large datasets. As sequencing costs continue to decline and large genome sequence datasets collected over the course of an outbreak become the norm, the need for a tractable solution to these problems grows [2]. Here we present the first steps towards such a solution by approximating, and then modifying, the model of [49].

In this manuscript we describe a novel birth-death-sampling model tailored for use in estimating the reproduction number and prevalence of infection in an epidemic. We start by reviewing existing sampling models for birth-death processes and derive a missing sampling model which has a natural interpretation in epidemiology, where data is usually only available in the form of binned (eg daily or weekly) counts. For example, if a health care provider is unable to report new cases over the weekend one might expect an aggregated number of cases to be reported at the start of the following week. This is in contrast to sequence data, which is often reported with the exact sampling date.

With several simulation studies we demonstrate empirically that our approximation (a) agrees with the output of an existing numerical scheme, (b) has linear complexity, considerably improving on existing computational approaches, which grow quadratically with the size of the data set, and (c) even with aggregated (binned) data, key parameters can still be recovered. Finally, we discuss the practical applications and benefits of TimTam and the limitations of our approach.

## Methods

Birth-death-sampling models are used to describe sequence data that have been either collected at predetermined points in time, hereafter *scheduled observations*, or opportunistically, ie when cases have presented themselves, hereafter *unscheduled observations* [29, 48]. The relationship between these sequences is described by the reconstructed phylogeny. The models of [51] and [49] consider an additional data type, which they term *occurrence data*, that represents unscheduled observation of infectious individuals without their inclusion in the reconstructed phylogeny. Such occurrence data may arise, for example, when an individual tests positive for infection but the pathogen genome is not sequenced.

We categorise observations based on two attributes, (i) whether the infected individuals were observed at predetermined times (scheduled observations) or follow a point process (unscheduled observations), and (ii) whether the observed cases were included in the reconstructed phylogeny (a *sequenced* observation), or not (an *unsequenced* observation).

This categorisation suggests an additional data type: the scheduled observation of unsequenced cases, which corresponds to the removal of multiple individuals from the infectious population at the same time, without incorporating them into the reconstructed phylogeny. There are several benefits to being able to incorporate such data. First, since epidemiological data are often given as a time series (instead of a point process) this is arguably a more natural way to utilise occurrence data in the estimation process [12]. The same could be said for the sequenced samples in instances when multiple samples are collected on the same day [27]. The second benefit is computational. Modelling observations as scheduled rather than unscheduled simplifies the likelihood, because a single scheduled observation can account for multiple unscheduled observations. As far as we are aware, scheduled unsequenced observations have not been considered in any phylodynamic inference method. Below we describe the sampling model formally and the method used to approximation of its likelihood, TimTam. An implementation of this method is available from (https://github.com/aezarebski/timtam).

### Phylogenetic Birth-Death Process

The birth-death (BD) process starts with a single infectious individual at the time of origin, *t* = 0. Infectious individuals “give birth” to new infectious individuals at rate *λ*, and are removed from the process either through naturally ceasing to be infectious (at rate *µ*, often called the “death” rate), or through being sampled. Unscheduled sampling of infectious individuals occurs at different rates depending on whether the samples are sequenced (which occurs at rate *ψ*) or not (which occurs at rate *ω*). An illustrative example of this process is shown in Panel A of Fig 1. Individuals can also be removed in scheduled sampling events. Scheduled sampling occurs at predetermined times, during which each infectious individual is independently sampled with a fixed probability: for a sequenced sample each lineages is sampled with probability *ρ* and for an unsequenced sample each lineage is sampled with probability *ν*. An illustrative example of the process with both scheduled and unscheduled sampling is shown in Fig S1. We denote scheduled sampling times *r*_*i*_ for sequenced sampling and *u*_*i*_ for unsequenced sampling, and assume these times are known a priori, since they are under the control of those observing the system.

**Fig 1.**
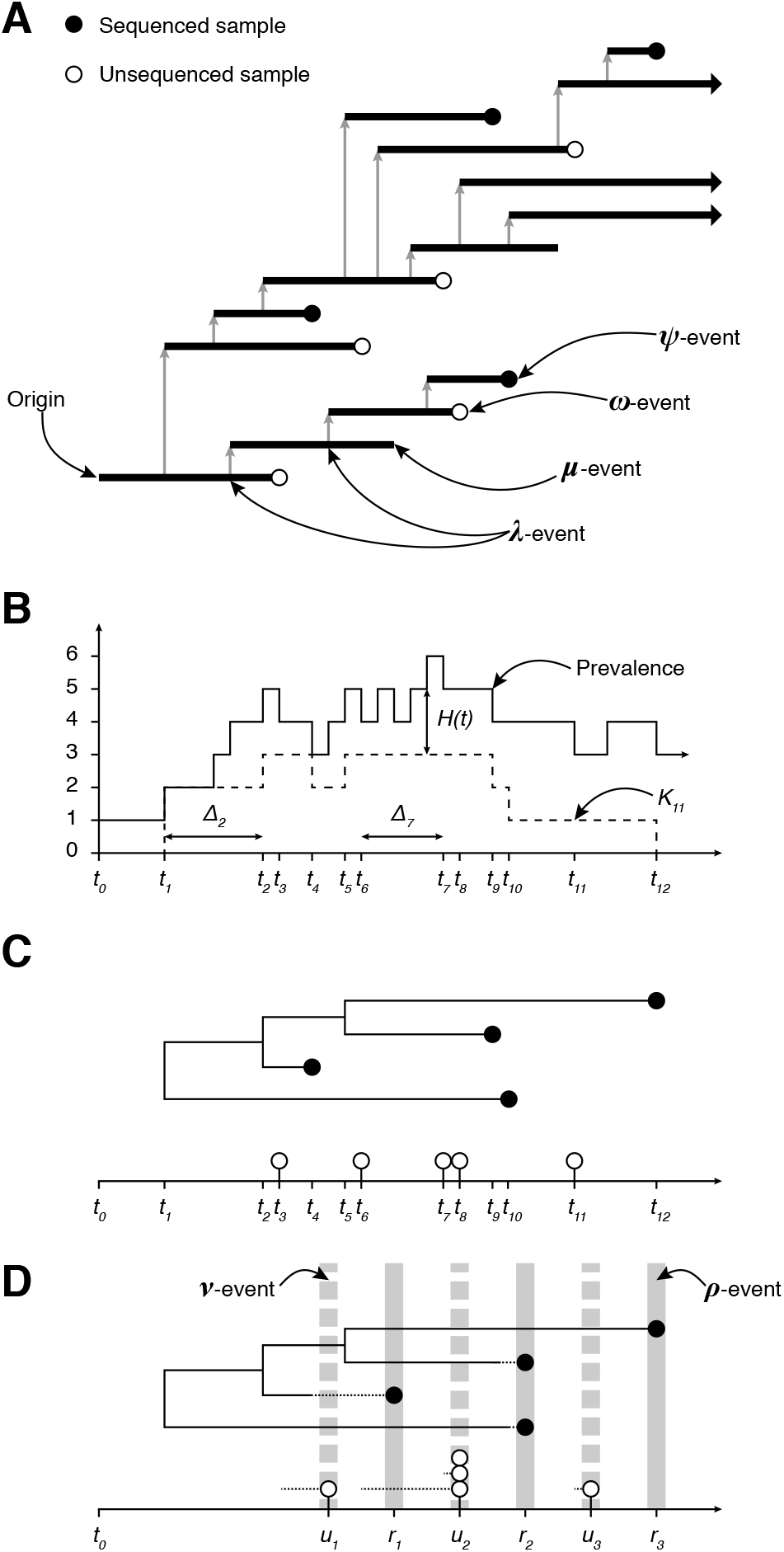
Birth-death model of transmission and observation. The process can be observed in several ways leading to different data types. **(A)** The transmission process produces a binary tree (the transmission tree) where an infection corresponds to a *λ*-event and a branch node and ceasing to be infectious corresponds to a *µ*-, *ψ*- or *ω*-event and a leaf node. **(B)** The number of lineages in the transmission tree through time, ie the prevalence of infection, and the number of lineages in the reconstructed tree, known as the lineages through time (LTT) plot, *K*.. **(C)** The tree reconstructed from the sequenced samples: *ψ*-events. The pathogen sequences allow the phylogeny connecting the infections and the timing of *λ*-events to be inferred. The unsequenced, *ω*-events form the point process on the horizontal axis. **(D)** Multiple *ψ*-events can be aggregated into a single *ρ*-event, such as the one at time *r*_2_. This loses information due to the discretization of the observation time, indicated by the dashed line segment. The same approach is used to aggregate *ω*-events into a single *ν*-event, eg the observation made at time *u*_2_.

Realisations of the process are binary trees with internal nodes corresponding to infection events and terminal nodes representing removal events as shown in Fig 1 and S1. We assume the edges of the tree are labelled with their length to ensure the nodes appear at the correct depth. The tree containing all infected individuals is the *transmission tree* (Fig 1A, and S1B). The subtree containing only the terminal nodes corresponding to sequenced samples (both scheduled and unscheduled) is called the *reconstructed tree* [39], (Fig 1C, and S1C). In practice, the topology and branch lengths of the reconstructed tree are estimated from the pathogen genomes; here we assume these are known a priori.

Trees can be summarised by *their lineages through time* (LTT) plot, which describes the number of lineages in the tree at each point in time. We denote the number of lineages in the reconstructed tree at time *t*_*i*_ by *K*_*i*_ (Fig 1B). We define the number of *hidden* lineages through time as the number of lineages that appear in the transmission tree but not in the reconstructed tree. The number of hidden lineages at time *t* is denoted *H*(*t*), and for convenience as *H*_*i*_ at time *t*_*i*_. The types of data that we consider can be thought of as a sequence of *N* events, *ε*_1:*N*_, starting from the origin and moving forward in time up to the present (ie the time of the last observation): *ε*_1:*N*_ = {(Δ*t*_*i*_, *e*_*i*_, Δ*K*_*i*_, Δ*H*_*i*_)}_*i*=1…*N*_ with Δ*t*_*i*_ denoting the time since the previous observation (ie Δ*t*_*i*_ := *t*_*i*_ − *t*_*i*−1_) and *e*_*i*_ describing the event that was observed at that time: *e*_*i*_ ∈ {*λ*-event, *ψ*-event, *ρ*-event, *ω*-event, *ν*-event}. The changes in the LTT and number of hidden lineages at time *t*_*i*_ are denoted Δ*K*_*i*_, so *K*_*i*_ = *K*_*i*−1_ − Δ*K*_*i*_, and Δ*H*_*i*_, so 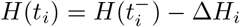.

There are two important assumptions in the description above. The first is that once and individual has been sampled they are removed from the infectious population. This is a standard, though not universal, assumption and often justified by the fact that sampling broadly coincides with receiving medical care, and hence taking care not to spread the infection further. The second is that if there is a scheduled sample, it contains either all sequenced samples or all unsequenced samples, ie there are no scheduled samples with both sequenced and unsequenced observations.

### The Likelihood

The joint conditional distribution of the process parameters, *θ* = (*λ, µ, ψ, ρ, ω, ν*), and the number of hidden lineages at time *t*_*N*_, *H*(*t*_*N*_), factorises as follows:

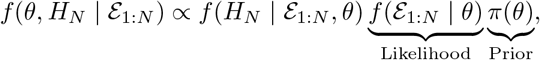

where *f* (*H*_*N*_ | *ε*_1:*N*_, *θ*) is the posterior distribution of the prevalence given *θ* which can be used to obtain the posterior predictive distribution of the prevalence: *f* (*H*_*N*_ | *ε*_1:*N*_). The likelihood has a natural factorisation which corresponds to processing the data from the origin through to the present:

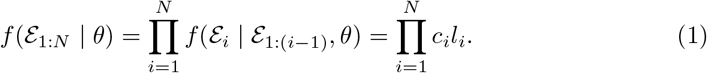

Since the likelihood of each observation depends on the distribution of the number of hidden lineages, the distribution of *ε*_*i*_ depends on the whole history *ε*_1:(*i*−1)_. Each factor, *f* (*ε*_*i*_ | *ε*_1:(*i*−1)_, *θ*), can be expressed as a product, *c*_*i*_*l*_*i*_, where *c*_*i*_ is the probability that no events where observed during the interval of time, (*t*_*i*−1_, *t*_*i*_), and *l*_*i*_ is the probability that the event observed at the end of the interval is *e*_*i*_.

Let *M* (*t, z*) be the generating function (GF) for the distribution of *H*(*t*) and the observations up until time *t*,

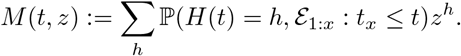

The likelihood is evaluated by traversing the data from the start of the process through to the present, calculating the distribution of hidden lineages and the *c*_*i*_ and *l*_*i*_ along the way.

Consider a sequence of functions, *M*_*i*_(*t, z*), which correspond to *M* (*t, z*) over the intervals (*t*_*i*_, *t*_*i*+1_), up to a normalisation constant which ensures *M*_*i*_(*t*_*i*_, 1) = 1. We define the *M*_*i*_ with a system of partial differential equations (PDEs) derived using the Master equations for the number of hidden lineages changes through time.

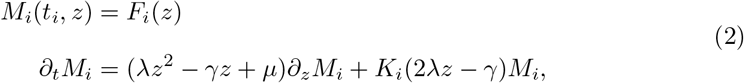

where *γ* = *λ* + *µ* + *ψ* + *ω* and *∂*_*x*_ is used to indicate partial differentiation with respect to the variable *x*. The number of lineages in the reconstructed tree, *K*_*i*_, only changes when there is a birth, or a sequenced sample and so is a constant over each interval.

The process starts with a single infected individual, so initially there are no hidden lineages and consequently the initial condition on the first interval is *M*_0_(0, *z*) = 1. Subsequent boundary conditions, *F*_*i*_(*z*), are based on the solution over the previous interval, *M*_*i*−1_ and the event that was observed at time *t*_*i*_.

The solution to Eq (2), first given as **Proposition 4.1** in [49], is

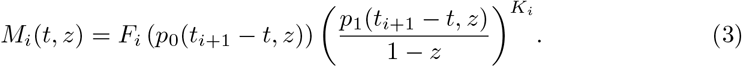

The functions *p*_0_ and *p*_1_ are standard results describing the probability of an individual and their descendents giving rise to exactly zero or one observation over a duration of length *t*_*i*+1_ − *t*; see [48] and the additional comments in the Appendix for further details.

Using Eq (3) the probability of not observing anything between times *t*_*i*_ and *t*_*i*+1_, and the probability generating function for the number of hidden lineages just prior to the observation at *t*_*i*+1_ are

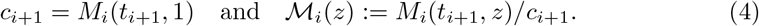

The process of calculating *l*_*i*+1_, the likelihood of observing *ε*_*i*+1_, and the next boundary condition, *F*_*i*+1_(*z*), the PGF of the number of hidden lineages at *t*_*i*+1_ is carried out in two steps. First, we transform *ℳ*_*i*_ to account for the observation of *ε*_*i*+1_ and evaluate the resulting expression at *z* = 1 to obtain *l*_*i*+1_ (using the transformations described below in Eq (5), (6), (7) and (8)). Second, we normalise the coefficients of this GF to get the PGF of *H*(*t*_*i*+1_), which is the boundary condition, *F*_*i*+1_(*z*), in the PDE for *M*_*i*+1_ in Eq (2). This process is repeated for each interval of time to get all the *c*_*i*_ and *l*_*i*_ in Eq (1).

We will now describe the transformations to *ℳ*_*i*_ used to account for the observation of *ε*_*i*+1_. Since *λ*- and *ψ*-events are only observed upon the reconstructed tree and do not influence the number of hidden lineages, *ℳ*_*i*_ is left unchanged when these are observed,

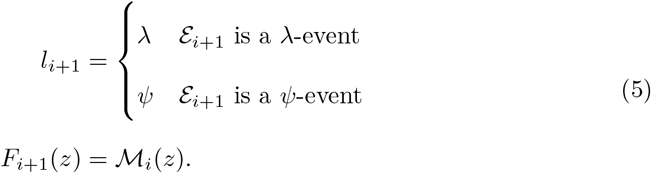

For an *ω*-event we need to shift the whole distribution of *H* and account for the unknown number of hidden lineages that could have been sampled, this is achieved by taking the partial derivative of the GF, which we denote by *∂*_*z*_, as elaborated upon in the Appendix. The likelihood of an *ω*-event is the normalising constant after the differentiation:

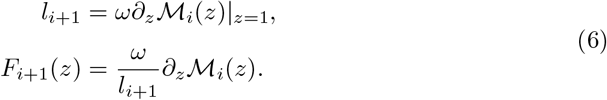

For a scheduled sampling event, at time *r*_*i*+1_ with removal probability *ρ*, we need to account for the survival of each of the *H*-lineages that were not sampled, those that were, and the number of lineages in the reconstructed tree that were not removed during this scheduled sampling. This leads to the following likelihood factor and updated PGF:

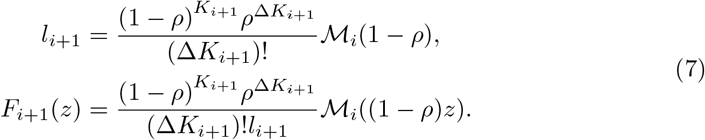

The factor of 1 − *ρ* in the argument of *ℳ*_*i*_ is to account for the *H*-lineages that were not sampled. The factors of 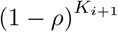 and 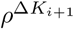 come from the lineages in the reconstructed tree that were not sampled (of which there are *K*_*i*+1_), and those that were sampled (of which there are Δ*K*_*i*+1_).

Last, we include scheduled unsequenced samples, ie the observation and simultaneous removal of multiple lineages without subsequent inclusion in the reconstructed phylogeny. For Equations (6), we noted that a single *ω*-sampling event corresponds to differentiating the PGF of *H* once. If at time *t*_*i*+1_ there is a scheduled unsequenced sample where each infectious individual is sampled with probability *ν*, and *n* lineages in total are sampled, then we must take the *n*-th derivative and accumulate a likelihood factor for the removed and non-removed lineages of (1 − *ν*)^*K*^*ν*^*n*^ (assuming the LTT at that time is *K*). We also have to scale *z* by a factor of 1 − *ν* to account for the *H*-lineages that were not sampled. Therefore, as in Equations (6) and (7), the likelihood and updated PGF after a *ν*-sample are:

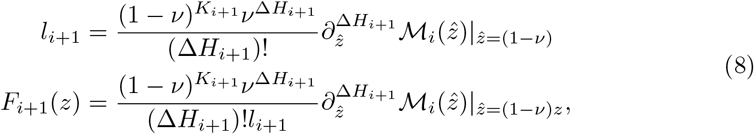

where the use of 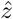 has been used to make explicit the order of operations.

Evaluating the expressions above numerically typically requires truncating a system of ordinary differential equations (ODEs) and solving them on each interval. This operation has a complexity which is cubic in the size of the truncated system (as a matrix exponential is required). [49] derives an approximation which has a quadratic complexity, albeit by introducing a further approximation. Our TimTam approximation, the main contribution of this paper, is as accurate as existing methods and has only a linear complexity.

### An Analytic Approximation

Our analytic approximation, TimTam, can be described as simply replacing the PGF of *H* with a more convenient PGF which describes a random variable with the same mean and variance. Specifically, we use the negative binomial (NB) distribution. We note two facts: first, we can evaluate the full PGF point-wise described above and, second, as shown in the Appendix, the GF of the negative binomial (NB) distribution is closed (up to a simple multiplicative factor) under partial derivatives and scaling of the parameter *z*. Together, these mean we can construct a NB approximation of the PGF at any point in the process and hence evaluate the resulting approximate likelihood and the distribution of hidden lineages. Algorithmically, this method can be expressed in the following steps:

1. Start at time *t*_*i*_ with the PGF *M*_*i*_ and use Equation (3) to obtain *M*_*i*_ at time *t*_*i*+1_.
2. Calculate *c*_*i*_ = *M*_*i*_(*t*_*i*+1_, 1^−^), the probability of not observing any events during the interval (*t*_*i*_, *t*_*i*+1_).
3. Define the PGF *ℳ*_*i*_ = *M*_*i*_*/c*_*i*_ and the PGF resulting from approximating it with a NB distribution: 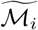.
4. Use 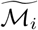 to compute, *l*_*i*_, the likelihood of observing *ε*_*i*+1_ and let *M*_*i*+1_ be the PGF of the number of *H*-lineages conditioning upon this observation (see Equations (6), (7) and (8).)
5. Increment the log-likelihood by log (*c*_*i*_*l*_*i*_) and return to Step 1 with an incremented *i* if there are remaining observations.

The steps involved require only the evaluation of closed form expressions and the number of iterations is linear with the number of observed events.

Our use of a NB moment-matching approximation is not arbitrary. [50] observed that the number of lineages descending from a single lineage has a zero-inflated geometric distribution and the sum of independent and identically distributed geometric random variables follows a NB distribution. Our approach of treating the number of lineages derived from *n* individuals as a NB random variable is somewhat motivated by combining these two properties. Further support for our approximation is obtained by considering an equivalent BD process, but with the modified total birth rate of *λn* + *a* where *a* is a small offset representing an immigration rate that leads to the removal of the extra (unobservable) zeros. Such processes can be described by NB lineage distributions at all times of their evolution and are stable to the inclusion of additional event types. [19, 24].

### Origin time vs TMRCA

The definition of the likelihood above assumes the origin of the phylogeny, *t*_0_ in Fig 1, is known or is a parameter to be estimated. This follows as we require the initial condition *M*_0_(0, *z*) = 1. In practice the phylogeny will likely only be known up to the time of the most recent common ancestor (TMRCA), *t*_1_ in Fig 1. We might account for this in one of two ways. The first, and simplest, is to treat the origin time as an additional parameter to be estimated. The second is to set a boundary condition at the TMRCA and to estimate the distribution of hidden lineages at that point, *H*_1_.

If one were confident the outbreak had stemmed from a single initial case, then the former method would be more suitable, especially if there was prior knowledge to constrain the time of origin. On the other hand, if we faced substantial uncertainty about how the outbreak began and sequencing was sparse, ie small *ψ* and *ρ*, then the TMRCA may be considerably more recent than the origin time and estimating the origin would be challenging. In this case, the latter approach may be more suitable. This would involve estimating the distribution of *H*_TMRCA_ and hence its GF *M*_1_(*t*_TMRCA_, *z*), from the family of NB distributions.

## Results

### Model validation and computational complexity

We performed a simulation study to compare TimTam with the method from [49], hereafter called the ODE approximation. The parameters used to generate a stratified set of simulations are given in Table 1. The S1 Appendix provides a full description of the simulation and subsampling process used to generate these test data. Fig 2 shows the value of the log-likelihood function evaluated using each method. Both methods produce very similar log-likelihood values, with TimTam explaining 98% of the variation in the ODE approximation values under a linear model.

**Table 1.** Parameters used for all simulated datasets.

| Parameter | Description | Value |
| --- | --- | --- |
| $\lambda$ | Birth rate | 1.7 |
| $\mu$ | Death rate | 0.9 |
| $\psi$ | Sequenced sampling rate | 0.05 |
| $\omega$ | Unsequenced sampling rate | 0.25 |
| $\rho$ | Scheduled sequenced sampling probability | 0.5 at $t = 6$ |

**Fig 2.**
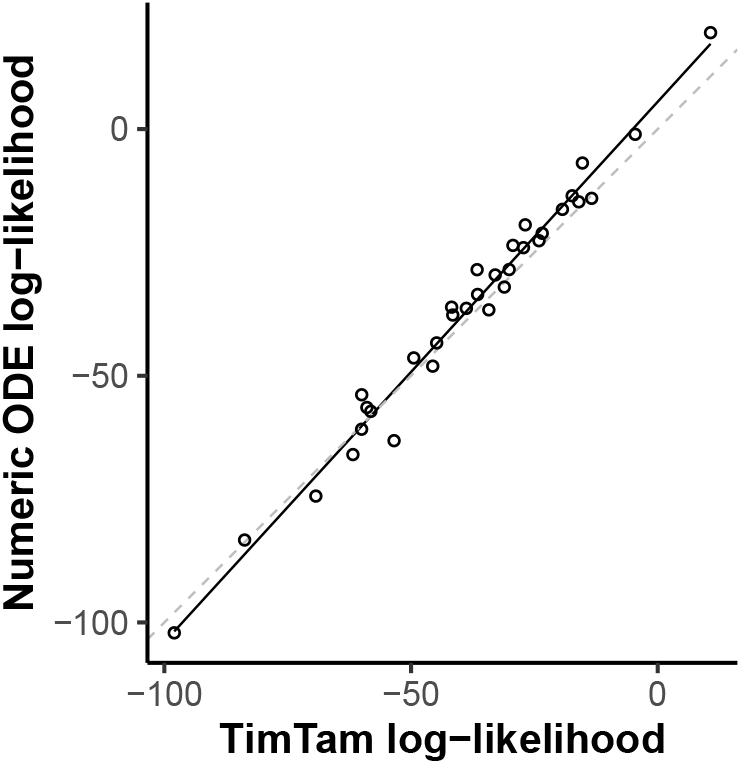
Likelihood comparison. Our TimTam approximation of the likelihood is in good agreement with the existing ODE approximation [49]. Each point shows the values of the log-likelihood computed using our approximation and the ODE approximation. The solid line shows a least squares fit which has an *R*^2^ of 0.98, the grey dashed line indicates parity, *y* = *x*.

To explore the computational complexity of TimTam, we measured how long it took to evaluate the log-likelihood for each of the simulated datasets. Fig 3 shows that with TimTam, the mean evaluation time grows approximately linearly with the size of the dataset, ∝ *n*^1.03^, where the 95% confidence interval (CI) on the exponent is (1.02, 1.04). In contrast, for the ODE approximation, the evaluation time grows approximately quadratically, ∝ *n*^2.38^, (95% CI = 2.26, 2.50). Since the ODE approximation requires specification of a truncation parameter, we obtained values for this parameter by increasing its value until doing so further resulted in a change to the log-likelihood of *<* 0.1%. The resulting truncation parameters are shown in Fig S2 in S1 Appendix. Full details of how the data were simulated, how the benchmarks were evaluated, and how the truncation parameter was selected are given in the Supplementary Materials.

**Fig 3.**
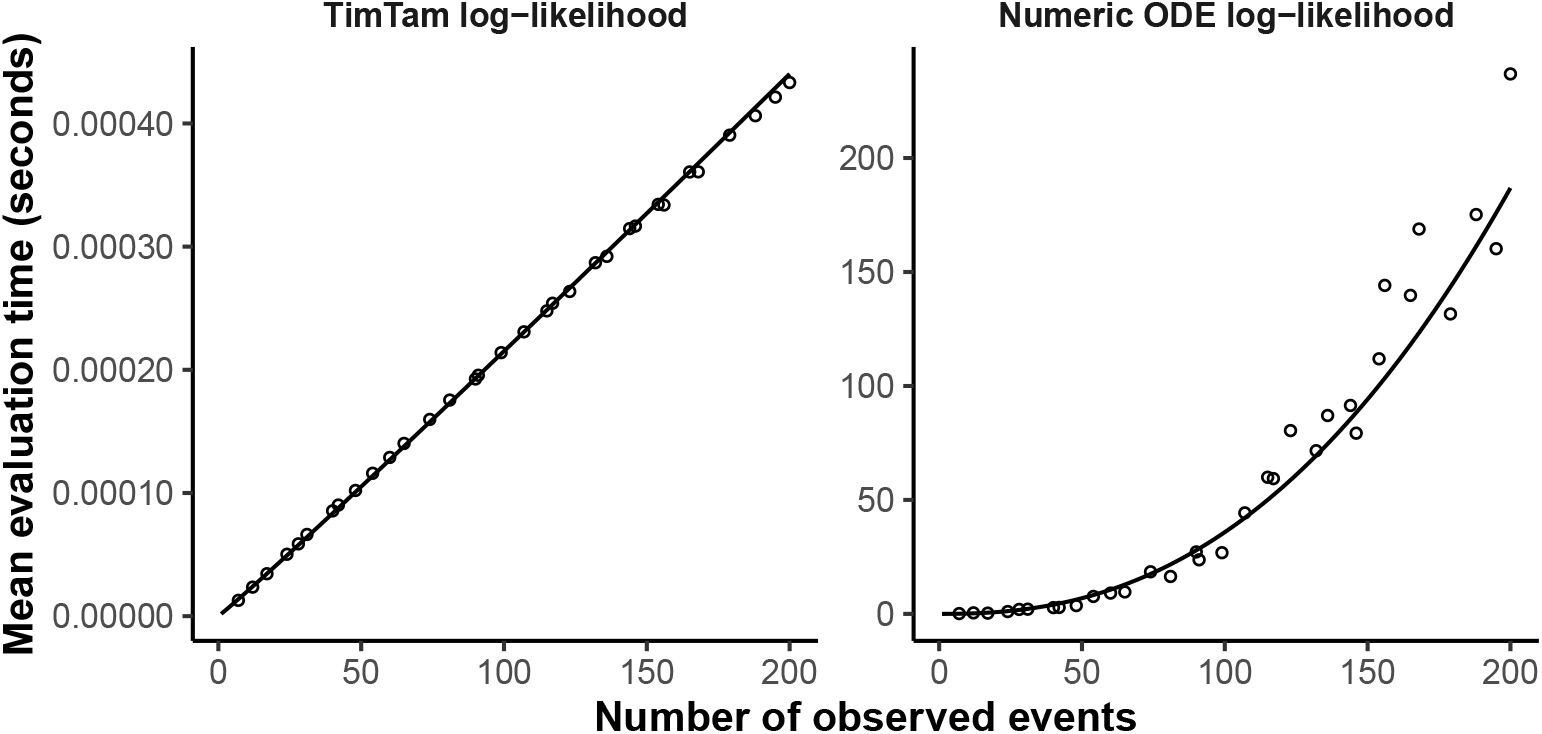
Log-likelihood evaluation time comparison. The time required to evaluate our approximation, TimTam, scales better with the dataset size than the existing ODE approximation. The scatter plots indicates the average number of seconds required to evaluate the log-likelihood function for each dataset size. The left panel contains the results using our approximation, which has times growing approximately linearly with the dataset size. The right panel contains the results using the ODE approximation, which has times growing approximately quadratically with the dataset size. Solid lines show least squares fits. Note that the *y*-axes are on different scales. The overall scaling factor (but not the exponent of the fitted model) may be implementation dependent.

In addition to the improvement in computational complexity, average evaluation times are orders of magnitude smaller for TimTam, which takes less than a millisecond in comparison to several seconds for the ODE approximation for larger datasets. However, we caution against over-interpreting the absolute computation times, since we used Haskell to implement TimTam, whereas the implementation of the ODE approximation, the same implementation used by [49], is a combination of C and Python. The faster computation time may depend on the programming language used as well as the algorithm. Nonetheless, the computational complexities of the respective algorithms means that the TimTam approach will outperform the ODE approximation for large datasets, regardless of the implementation.

### Parameter identifiability and aggregation scheme

Having validated TimTam against the ODE approximation, we now showcase our approach as an estimation scheme that merges all the data types considered in this manuscript. We also explore the effect of aggregating unscheduled samples into scheduled sampling events, looking at the accuracy and bias of the estimates when we further obfuscate the data.

We first verified that, given a known death rate *µ*, the model parameters are identifiable using a simulation that includes all four types of sampling events described above. Fig S3–S9 of S1 Appendix show cross sections of the likelihood surface and scatter plots of the posterior samples. We also show that the statistical power to estimate model parameters increases with simulation length (and hence the size of the dataset). Additional details of the simulation and estimation methods are given in S1 Appendix.

Next, we simulated a dataset using the rate parameters in Table 1 but with the scheduled sampling probability set to zero, ie a simulation which only contains unscheduled samples. The simulation was started with a single infectious individual and stopped at *t* = 13.5. From the unscheduled observations a second dataset was derived, this was done by aggregating the unscheduled observations into scheduled observations, eg all the unscheduled sequences sampled during the interval (*t*_*a*_, *t*_*b*_] were combined into a single scheduled sequenced sample at time *t*_*b*_ (as illustrated in Fig 1D). This aggregation reflects how cases may only be reported at particular temporal resolutions, eg daily or weekly case counts.

The sequenced samples were aggregated into observations at *t* = 2.5, 3.5, …, 13.5 and unsequenced samples were aggregated at *t* = 2.4, 3.4, …, 13.4. In simulating these data, only simulations that did not go extinct during the simulation period and had 1000–10000 events were used (as a way to avoid excessive run times and ensure that there was a sufficient amount of transmission). Moreover, any simulations where the simulated population decreased to only a single individual at any time after the first infection were discarded, as this could result in the reconstructed tree having a significantly younger TMRCA than the transmission tree.

Fig 4A and B shows the sequenced and unsequenced samples in the simulated dataset. Fig 4C and D shows the same dataset after aggregation. Fig 4E shows the prevalence through time in the simulation and the corresponding estimates at *t* = 13.5 using the simulated and aggregated datasets, respectively. Fig 5 shows the marginal posterior distributions of *λ*, and either *ψ* and *ω*, or *ρ* and *ν* depending on the dataset used.

**Fig 4.**
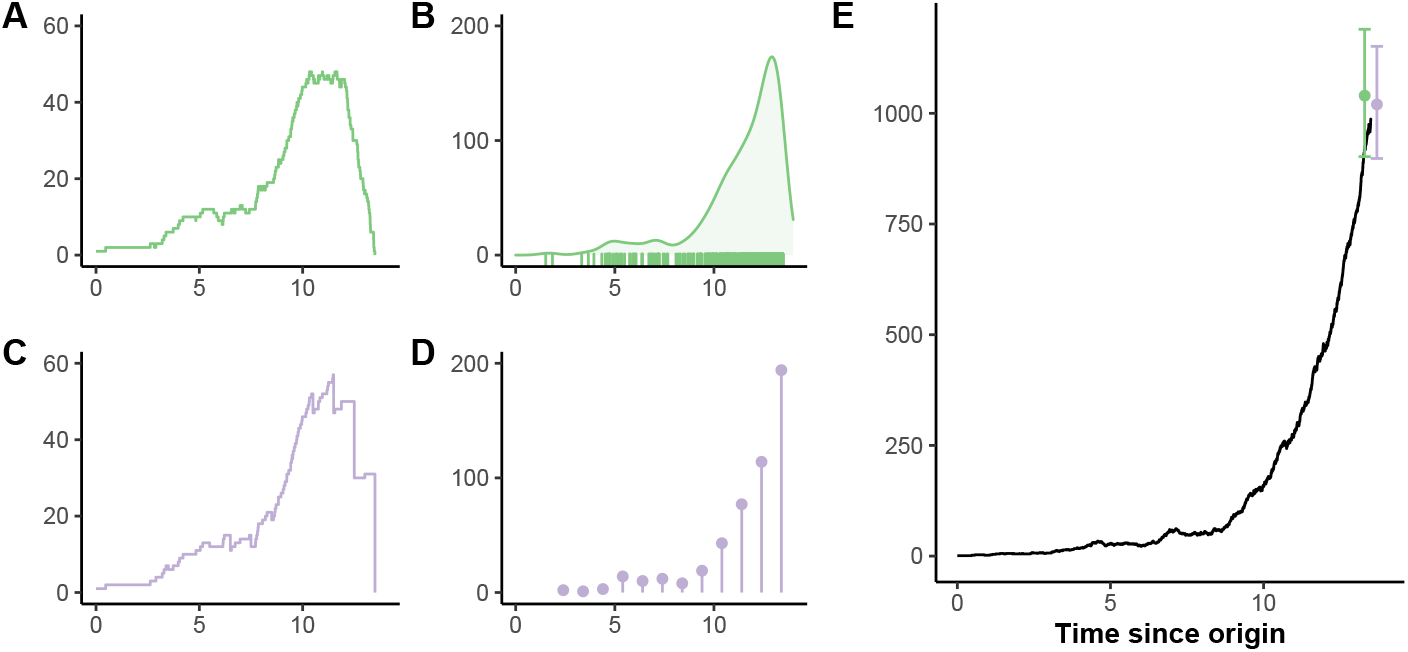
Data aggregation example. The effect of aggregation on the dataset and estimates of prevalence. **(A)** The LTT of the tree reconstructed from the unscheduled sequenced observations. **(B)** The density of unscheduled unsequenced observations, ie a point process of observations. **(C)** The LTT of the tree reconstructed from the sequenced observations after aggregation into scheduled sampling events. **(D)** The number of unsequenced observations aggregated into regular scheduled observations, ie a time series of cases reported at regular intervals. **(E)** The total prevalence of infection throughout the simulation is represented by the black line, the points and error bars indicate estimates (and 95% credible intervals) of the prevalence at the present, colour coded by the dataset used (green, unscheduled data; lilac, aggregated data). Fig 5 shows the marginal posterior distributions using each dataset.

**Fig 5.**
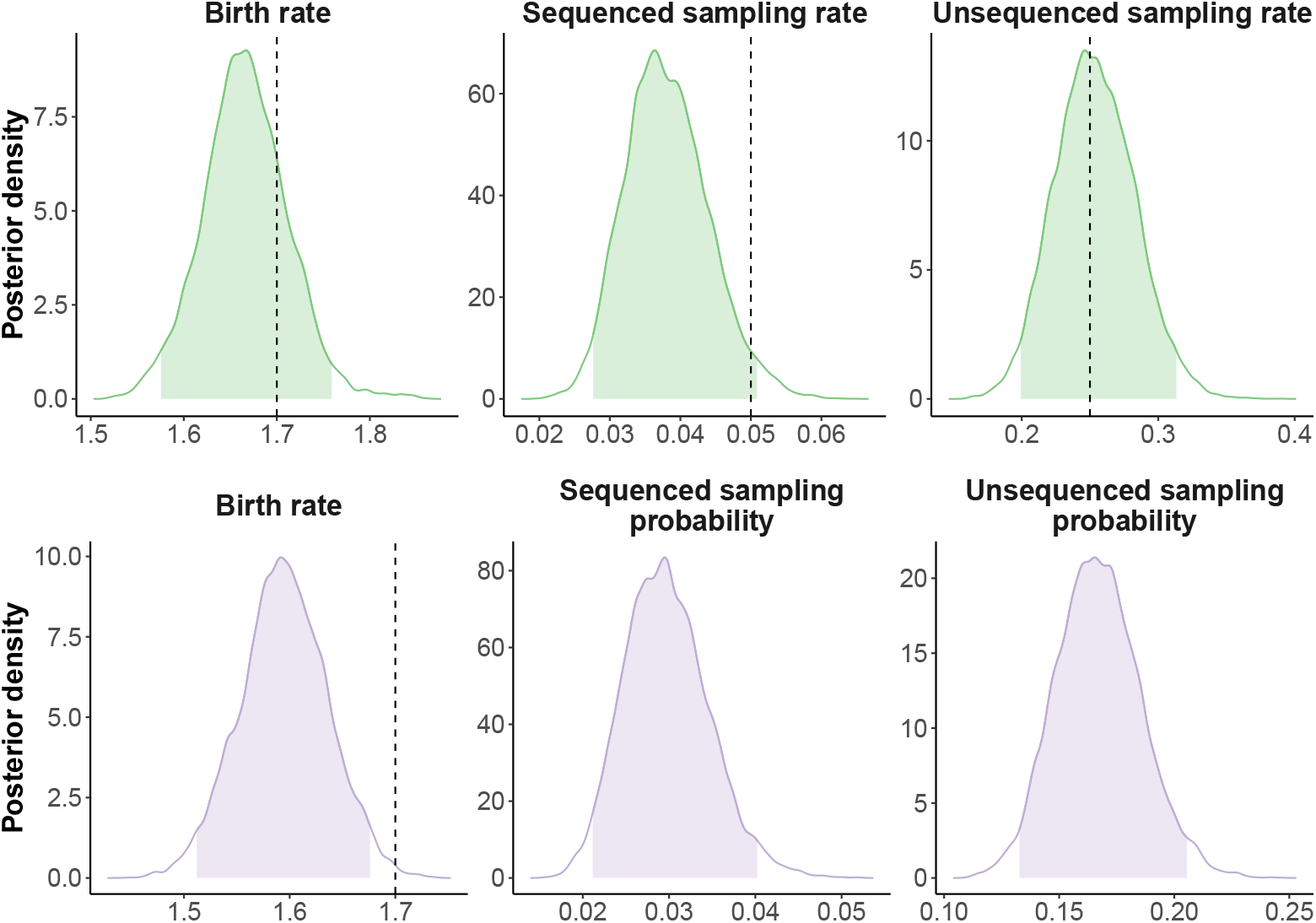
Posterior distributions. Given the death rate, *µ*, the posterior distributions for both datasets shown in Fig 4 have well-defined maxima. The charts show the marginal posterior distributions of parameters using either the unscheduled samples (top row, green) or the scheduled samples post aggregation (bottom row, lilac). Filled areas indicate 95% credible intervals. Vertical dashed lines indicate true parameter values where they exist (Table 1). There are no vertical lines for the scheduled observation probabilities because they are not well defined for this simulation.

When estimating model parameters the death rate *µ* was fixed to the true value used while simulating the data, since not fixing one of the parameters makes the likelihood unidentifiable and estimates of *µ* may be obtained from additional data sources [4, 29]. The posterior samples where generated via MCMC. Standard diagnostics were used to test the convergence and mixing of the MCMC, (further details of the MCMC diagnostics and visualisations of the joint distribution of the posterior samples are given in S1 Appendix.)

While prevalence estimates from both the original unscheduled and aggregated datasets are overlapping and contain the truth, aggregation leads to underestimating the birth rate. This bias is likely due to the aggregation scheme used (see S1 Appendix for further commentary). Moreover, the sequenced sampling rate is underestimated when using the unscheduled dataset. We conjecture that this is due to there being roughly five times fewer sequenced than unsequenced samples. Although the true values for the sampling probabilities estimated from the aggregated dataset are not known, the ratio between the two parameters is similar to the ratio between the unscheduled sampling rates.

### Repeated simulation to test credible interval coverage

Fig 6 (top panel) shows the 95% credible interval (CI) and point estimate (posterior median) of the basic reproduction number, ℛ_0_ = *λ/*(*µ* + *ψ* + *ω*), for each of 100 simulation replicates. The simulation parameters used are the same as those used to simulate the data shown in Fig 4. The estimates are sorted according to the estimated ℛ_0_ value. Of the 100 replicates, 87 have a CI containing the true ℛ_0_. The Appendix contains some commentary on the level of coverage that is expected.

**Fig 6.**
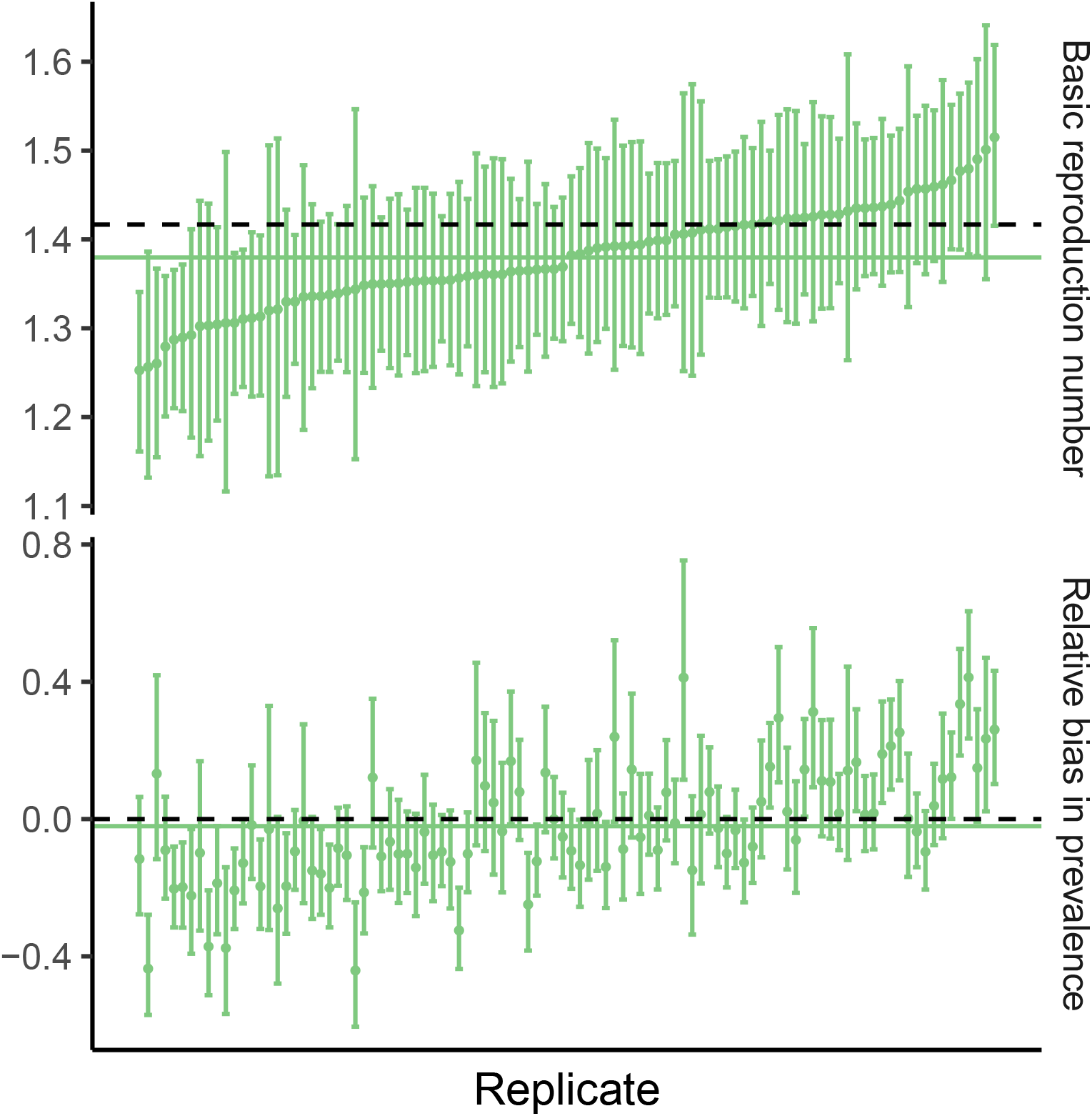
Simulation study results. The bias in the estimators of the basic reproduction number, ℛ_0_, and the prevalence is small. The top panel shows the (ranked) ℛ_0_ point estimates and 95% CI for each replicate. For 87 of these the CI contains the value used in the simulation, 1.42, which is indicated by the horizontal dashed line. The bottom panel shows the relative error in the prevalence estimate (ie a value of zero corresponds to the true prevalence in that replicate.) The coverage (64 of 100) is lower than 95% which is not unusual given coverage properties do not hold in general for credible intervals. The corresponding intervals using the aggregated data are shown in Figures S8 and S9. The solid horizontal lines indicate the mean of the point estimates.

Fig 6 (bottom panel) shows the 95% CI and point estimate (posterior median) of the relative bias in the estimate of the prevalence in each replicate (ie the proportion by which the estimate differs from the true prevalence in that particular replicate; for an estimate 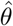 of *θ*, this is 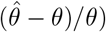. The relative bias is used rather than the bias because the true prevalence varies substantially across replicates making it difficult to compare them. In this figure the replicates in the top and bottom panels are in the same order. Of the 100 replicates, 64 have a CI containing the true prevalence at the end of the simulation (and hence cross 0).

Analogous estimates were performed for the aggregated data (generated using the process described above). It appears that the aggregation introduces a systematic bias towards underestimation of the birth rate. The estimates of the prevalence at the present are similarly unbiased for the aggregated data, although the CI coverage is lower. Full results are presented in S1 Appendix.

## Discussion

We have described an analytic approximation, called TimTam, for the likelihood of a birth-death-sampling model which can also describe *scheduled data* ie cohort sampling or reporting at predetermined times. TimTam can analyse both sequenced and unsequenced samples, ie the observations can represent sequences that are either included in the reconstructed tree, or observed infections that are not sequenced (occurrence data). Our approach generalises previous birth-death estimation frameworks [47, 49, 51] by accommodating and exploiting more data types than previously considered and makes it feasible to analyse very large datasets.

Our work is a step towards more flexible time series-based approaches to phylodynamics, in which multiple sequences are processed concurrently as elements of a time series. This extends the more common point-process based paradigm, in which samples are considered individually. TimTam also provides an estimate of the distribution of the prevalence of infection, allowing both the estimation of summary statistics, such as ℛ_0_, and the total number of cases. Comparison with existing algorithms on small-to-moderate sized datasets suggests it faithfully represents the true likelihood function.

At present, we cannot provide rigorous bounds on the error introduced by this approximation (although work is underway on this). Based on our simulation study, the credible intervals under this likelihood (with an improper uniform prior) slightly underestimate the level of uncertainty in the estimates of the basic reproduction number and the prevalence of infection. Although, as discussed, this is not surprising given these are credible intervals rather than confidence intervals.

Based on work from [50], we conjecture that if the probability of extinction becomes large, the zero inflation in the geometric distributions describing the number of descending lineages might become an issue. Since our focus is on large datasets describing established epidemics, we expect that this situation will rarely arise in practice. Additionally, as the death rate increases, the power of birth-death models as an inference tool is naturally limited by a lack of data [35, 36]. If this method is applied to small outbreaks or, when the reproduction number is low, sensitivity analyses will be necessary to check the fidelity of the negative binomial approximation.

Our work echoes the frameworks of [51] and [49], but trades some generality for simplicity and tractability. Specifically, [51] presented a particle filtering method that can be applied more generally, while [49] derived a complete posterior predictive distribution of prevalence over time, which allows the study of historical transmission. Another limitation of our approach, which is common to many models, is to neglect *sampled ancestors*, ie individuals who have been observed but remain in the infectious population [47, 49, 54]. While the former can describe a greater variety of birth-death processes and the latter can be used to estimate additional properties of the process, the scalability of both frameworks are limited by the computational burden.

Our approximation provides a computationally efficient method for handling diverse data types (such as data aggregated to a daily or weekly resolution) that is scalable to large datasets. We also introduce an aggregation scheme that radically reduces the computational burden with only a modest expense to the accuracy. The improvement in performance stems from the resulting likelihood computation scaling by the number of aggregated intervals, proportional to epidemic length, rather than the epidemic size. In many real epidemic scenarios data are only reported at a particular temporal resolution and in such scenarios this aggregation reflects the best-case for inference. As the availability of phylogenetic data (derived from sequences or contact-tracing) increases and the size of these data grows, such approximation schemes will become increasingly valuable.

## Supporting information

**S1 Appendix. Additional details of the approximation scheme and computational methodology**. This document provides additional details regarding the derivation of the approximation scheme and provides additional detail on the simulation and benchmarking computations.

## Acknowledgements

The authors are grateful to Marc Manceau for his keen examination of an earlier draft of this manuscript and helpful comments.

AEZ, OGP and LdP are supported by The Oxford Martin Programme on Pandemic Genomics. KVP is funded under grant reference MR/R015600/1 by the UK Medical Research Council (MRC) and the UK Department for International Development (DFID)

