## Appendix for "A computationally tractable birth-death model that combines phylogenetic and epidemiological data"

> A generating function is a clothesline on which we hang up a sequence of numbers for display.
>
> Herbert S. Wilf, *Generatingfunctionology*

#### Theoretical results

##### Probability Generating Functions (PGF)

Recall that for a random variable, $X$, its probability generating function (PGF) in the variable $z$ is

$$G_X(z) := \mathbb{E}_X\left[z^X\right].$$

Some useful elementary properties of $G_X$ are that:

- $G_X(1^-) = 1$.
- $G'_X(1^-) = \mu$ where $\mu$ is the expected value of $X$.
- $\sigma^2 = G''_X(1^-) + G'_X(1^-) - (G'_X(1^-))^2$ where $\sigma^2$ is the variance of $X$.
- $G_S(z) = G_N(G_X(z))$ if $S = \sum_i^N X_i$ for IID $X_i$ and has $G_{X_1+\cdots+X_n}(z) = G_X(z)^n$ as a special case.

Consequently, a generating function $H(z)$ which has positive coefficients with a finite sum, induces a distribution with PGF $G(z) := H(z)/H(1^-)$. In this case, we refer to $H(1^-)$ as the normalising constant.

##### Properties of the negative binomial distribution

Consider a negative binomially (NB) distributed random variable, $X \sim \text{NegBinom}(r, p)$. Its probability mass function (PMF) is:

$$\mathbb{P}(X = n) = \binom{n+r-1}{n}(1-p)^r p^n$$

The parameters $r$ and $p$ can be expressed in terms of the mean, $\mu$, and variance, $\sigma^2$, as:

$$p = \frac{\sigma^2 - \mu}{\sigma^2} \quad \text{and} \quad r = \frac{\mu^2}{\sigma^2 - \mu}. \tag{1}$$

The probability generating function (PGF), $G(z; p, r)$ for this random variable is:

$$G(z; p, r) = \left(\frac{1-p}{1-pz}\right)^r \text{ where}$$
$$p = \frac{\sigma^2 - \mu}{\sigma^2} \quad \text{and} \quad r = \frac{\mu^2}{\sigma^2 - \mu}. \tag{2}$$

It is stated in the Methods section of the main text, if $H$ has a negative binomial distribution, then conditioning on each of the observations leaves it with a negative binomial distribution. To see this, note both $\lambda$- and $\psi$-events do not influence $H$, hence conditioning upon them does not alter the distribution of $H$. Moreover, as the family of negative binomial PGFs is closed (up to a multiplicative constant) under both scaling of $z$ and partial derivatives with respect to $z$, it can be shown by induction that:

$$\partial_z^n G(z; p, r) = r^{\bar{n}} \left(\frac{p}{1-p}\right)^n G(z; p, r+n),$$

where $x^{\bar{n}} := x(x+1)\ldots(x+n-1)$ is the rising factorial (a.k.a. the Pochhammer function). The corresponding result for scaling $z$ is the following:

$$\begin{aligned} G(\alpha z; p, r) &= \left(\frac{1-p}{1-p\alpha}\right)^r \left(\frac{1-p\alpha}{1-p\alpha z}\right)^r \\ &= \left(\frac{1-p}{1-p\alpha}\right)^r G(z; p\alpha, r). \end{aligned}$$

These results are important for conditioning the distribution of $H$ on both $\rho$- and $\omega$-events.

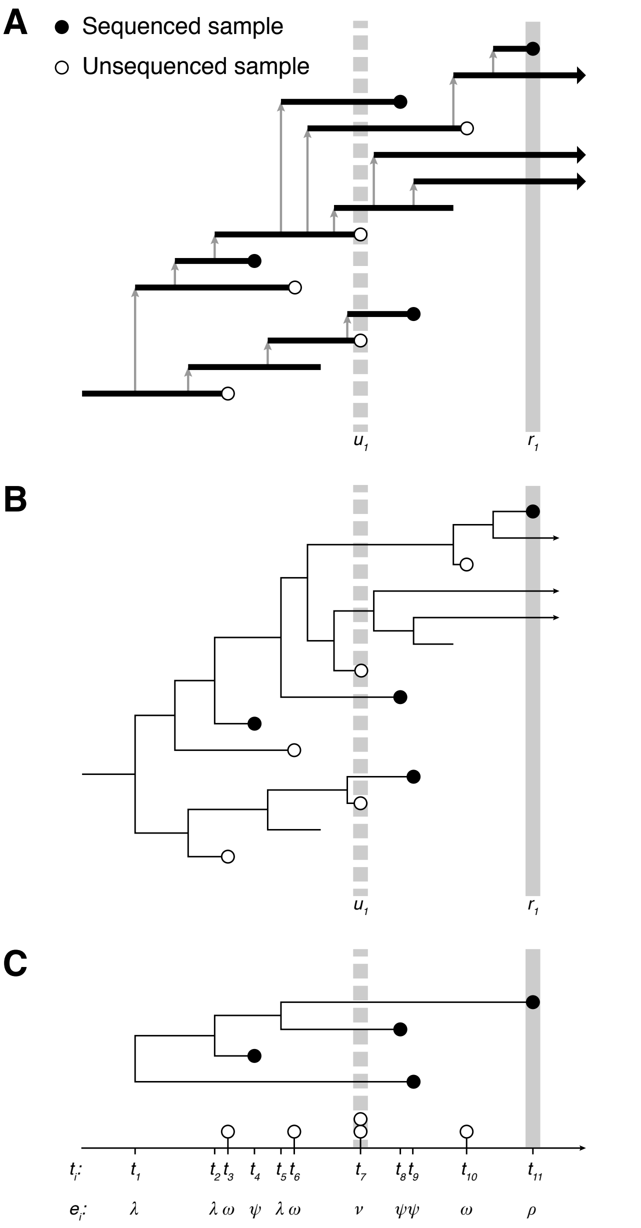


**Figure S1: Birth-death model of transmission and observation with scheduled samples.** In addition to unscheduled sampling which occurs continuously, we consider scheduled sampling where at predetermined times a binomial sample of the infectious population is removed. This corresponds to a cross-sectional study of prevalence. **(A)** The vertical lines indicate the timing of the scheduled samples: the dashed line (at time $t_7$) is an unsequenced sample which observed two infectious individuals, the solid line (at time $t_{11}$) is a sequenced sample. **(B)** The transmission tree corresponding to the realisation of the birth-death process, which appears in Panel A. **(C)** The reconstructed tree with sequenced observations on its leaves and the unsequenced observations as a point process. The example in this figure differs from Fig 1 of the main text in that here none of the unscheduled samples have been aggregated, the scheduled data has been generated as part of the observation process.

#### Useful results for birth-death processes

Here we describe some results from the existing literature modified to match the notation and model we have adopted in the current work[1]. First we have a couple of results adapted from **Theorem 3.1** of [Sta10]. Consider the birth-death process (and notation) described in the Methods section of the main text, the probability that an individual, alive at time $t$, will generate no $\psi$-, $\omega$- or $\rho$-sampled observations by time $T$ when there is a $\rho$-sampling event. Let $p_0(u)$ denote this probability where $u := T - t$, then $p_0$ must satisfy the following differential equation:

$$p_0(0) = z \quad \text{and} \quad \frac{dp_0}{du} = \mu - \gamma p_0(u) + \lambda p_0(u)^2,$$

where the value of $z$ will typically be $1 - \rho$ to denote the probability that the lineage was not $\rho$-sampled at time $T$. The solution is:

$$p_0(u, z) = \frac{x_1(x_2 - z) - x_2(x_1 - z)e^{-\sqrt{\Delta}u}}{(x_2 - z) - (x_1 - z)e^{-\sqrt{\Delta}u}} \tag{3}$$

where

$$x_1 = \frac{\gamma - \sqrt{\Delta}}{2\lambda}, \quad x_2 = \frac{\gamma + \sqrt{\Delta}}{2\lambda}, \\ \gamma = \lambda + \mu + \psi + \omega \quad \text{and} \quad \Delta = \gamma^2 - 4\lambda\mu. \tag{4}$$

In a similar manner, the probability of there being exactly one $\rho$-sampled lineage and no sampled extinct lineages, the function $p_1(u)$, satisfies the differential equation:

$$p_1(0) = 1 - z \quad \text{and} \\ \frac{dp_1}{du} = -\gamma p_1(u) + 2\lambda p_0(u)p_1(u). \tag{5}$$

Note that for this equation, the initial condition is $1 - z$ since it will typically be used to indicate that the lineage was $\rho$-sampled at time $T$. Using the definitions in Equation (4), the solution is:

$$p_1(u, t) = \frac{(1 - z)\Delta}{\lambda^2} \frac{e^{-\sqrt{\Delta}u}}{((x_2 - z) - (x_1 - z)e^{-\sqrt{\Delta}u})^2}. \tag{6}$$

These results were used in [MGVS20] to derive the generating function, $M(t, z)$, for the number of lineages that do not appear in a phylogeny during an interval of time without any observed events. The following result is adapted from **Proposition 4.1** of [MGVS20]. Consider the process described in the Methods section of the main text, during a period of time $[a, b]$ during which there are $k$ lineages and no observed events, if the probability generating function for the number of lineages which do not appear in the phylogeny is initially $F(z)$, then the generating function satisfies the following PDE:

[1] Note that care needs to be taken interpreting the following results due to the reversal of time. Since the current manuscript is written in terms of forward time we have tried to remain consistent in this notation.

$$
\begin{aligned}
&M(a,z) = F(z) \quad \text{and} \\
&\quad \partial_t M = (\mu - \gamma z + \lambda z^2)\partial_z M + k(2\lambda z - \gamma)M
\end{aligned} \tag{7}
$$

The solution to this is:

$$M(t,z) = F(p_0(b-t,z))\left(\frac{p_1(b-t,z)}{1-z}\right)^k.$$

#### Unsequenced samples correspond to partial derivatives

First, we consider the case of *unscheduled*, unsequenced samples, which remove one $H$-lineage and occur with rate $\omega$. Let $\mathcal{M}(z)$ be the generating function for the number of $H$-lineages prior to an observation. The $j$-th term of this series is $h_j z^j$, where $h_j$ is the probability that the number of $H$-lineages is $j$. We want to find the corresponding term after one of the lineages has been removed at random. Since there are $j$ lineages, there are $j$ ways to sample one lineage, and upon sampling it is removed from the population (which then only has $j-1$ lineages.) Therefore, the term $h_j z^j$ becomes $\omega j h_j z^{j-1}$ (ie it forms the $(j-1)$-th term of the resulting generating function). Summing over $j$ we find that the resulting generating function is equal to $\omega\partial_z\mathcal{M}(z)$. Selecting one of the lineages corresponds to the operation of *pointing* in combinatorics [FS09], which would mean we take the partial derivative, $\partial_z$ and then multiply by $z$. However, since we remove the lineage after selecting it, the results differ by a factor of $z$

For *scheduled* unsequenced samples, each $H$-lineage is sampled (and removed) with probability $\nu$, or remains with probability $1-\nu$. Consider the case where $\Delta H$ of the $H$-lineages have been sampled. If there were $j$ lineages to start with, the probability of sampling $\Delta H$ is:

$$
\begin{aligned}
&\binom{j}{\Delta H}\nu^{\Delta H}(1-\nu)^{j-\Delta H} \\
&= \frac{1}{(\Delta H)!}(j)_{\Delta H}\nu^{\Delta H}(1-\nu)^{j-\Delta H},
\end{aligned}
$$

where $(x)_n = x(x-1)\dots(x-n+1)$ is the Pochhammer symbol. As above, we can then write down the terms of the generating function after the scheduled sampling event and sum over $j$ to find the new generating function,

$$\frac{\nu^{\Delta H}}{(\Delta H)!}\partial_z^{\Delta H}\mathcal{M}((1-\nu)z)$$

where $\partial_z^n$ indicates the $n$-th partial derivative.

#### Statistics of $H$ via the generating functions $M_i$

The following equations describe the application of the properties above to the generating function for $H$. Consider the partial derivatives of $M_i$ (which become relevant in the next section). Let $A = x_2 - x_1 e^{-\sqrt{\Delta}u}$, $B = 1 - e^{-\sqrt{\Delta}u}$ and $C = x_2 e^{-\sqrt{\Delta}u} - x_1$ in the expression for $p_0$ from Equation (3), we get the following form which simplifies some subsequent calculus:

$$
\begin{aligned}
p_0(u,z) &= \frac{x_1(x_2-z) - x_2(x_1-z)e^{-\sqrt{\Delta}u}}{(x_2-z)-(x_1-z)e^{-\sqrt{\Delta}u}} \\
&= \frac{x_1x_2(1-e^{-\sqrt{\Delta}u}) + (x_2e^{-\sqrt{\Delta}u} - x_1)z}{(x_2 - x_1e^{-\sqrt{\Delta}u}) - (1-e^{-\sqrt{\Delta}u})z} \\
&= \frac{x_1x_2B + Cz}{A - Bz}
\end{aligned} \tag{8}
$$

The expression for $p_1$ from Equation (6) can also be expressed in terms of $A$ and $B$ in a convenient form:

$$
\begin{aligned}
\frac{p_1(u,z)}{1-z} &= \frac{\Delta}{\lambda^2}\frac{e^{-\sqrt{\Delta}u}}{((x_2-z)-(x_1-z)e^{-\sqrt{\Delta}u})^2} \\
&= \frac{\Delta e^{-\sqrt{\Delta}u}}{\lambda^2}\frac{1}{(A-Bz)^2},
\end{aligned} \tag{9}
$$

In subsequent calculations we will need the partial derivatives (with respect to $z$) for both $p_0$ and $p_1/(1-z)$ which are:

$$
\begin{aligned}
\partial_z p_0(u,z) &= \frac{CA + x_1x_2B^2}{(A-Bz)^2} \quad \text{and} \\
\partial_z^2 p_0(u,z) &= \frac{2B(CA + x_1x_2B^2)}{(A-Bz)^3}.
\end{aligned} \tag{10}
$$

and

$$
\begin{aligned}
\partial_z\left(\frac{p_1(u,z)}{1-z}\right) &= \frac{\Delta e^{-\sqrt{\Delta}u}}{\lambda^2}\frac{2B}{(A-Bz)^3} \quad \text{and} \\
\partial_z^2\left(\frac{p_1(u,z)}{1-z}\right) &= \frac{\Delta e^{-\sqrt{\Delta}u}}{\lambda^2}\frac{6B^2}{(A-Bz)^4}.
\end{aligned} \tag{11}
$$

We also have the following:

$$
\begin{aligned}
M(u,z) =& F(p_0(u,z))\Bigg(\underbrace{\frac{p_1(u,z)}{1-z}}_{R(u,z)}\Bigg)^k \\
\partial_z M(u,z) =& F'(p_0(u,t))\partial_z p_0(u,z)R(u,z)^k + \\
& F(p_0(u,z))kR(u,z)^{k-1}\partial_z R(u,z) \\
\partial_z^2 M(u,z) =& \\
& F''(p_0(u,z))(\partial_z p_0(u,z))^2 R(u,z)^k + \\
& F'(p_0(u,z))(\partial_z^2 p_0(u,z))R(u,z)^k + \\
& 2F'(p_0(u,z))(\partial_z p_0(u,z))kR(u,z)^{k-1}\partial_z R(u,z) + \\
& F(p_0(u,z))k(k-1)R(u,z)^{k-2}(\partial_z R(u,z))^2 + \\
& F(p_0(u,z))kR(u,z)^{k-1}\partial_z^2 R(u,z)
\end{aligned} \tag{12}
$$

Although Equation (12) appears messy, the number of expressions that need to be evaluated can be reduced using Equations 10 and 11.

### Computational results

#### Simulation and selection of truncation parameter

To compare the computational cost of evaluating the ODE approximation [MGVS20] with our TimTam approach, we simulated datasets of varying size and measured the time it took to evaluate the log-likelihood for these datasets using each algorithm. This also demonstrates the degree to which the two approximations agree on the value of the log-likelihood.

We simulated 1000 realisations of the birth-death process using the parameters shown in Table 1 of the main text. Each simulation was started with a single infectious individual and terminated at time $t = 6$, at which point there is a scheduled sequenced sampling event with probability $\rho = 0.5$. These simulations where then filtered to get a more uniform distribution of dataset sizes. This was done by selecting the first simulation which contained a number of events which fell in a variety of ranges: 1–5, 6–10, etc, up to 196–200. Any intervals that did not contain a simulation with one of these sizes was left empty. This filtering process left 33 simulated data sets.

The ODE approximation has a truncation parameter which needs to be set to a large value but for which no selection criterion has been provided. To select the truncation parameter, starting from a value of 15 we increased it in increments of 5 until a value was reached where an additional increment did not change the value of the log-likelihood by more than 0.1%. If no such value was reached below a threshold of 125 then this simuation was removed from the test set. The resulting truncation parameter selected for each simulation is shown in Figure S2. The selected truncation parameter tends to grow linearly with the number of observations in the dataset.

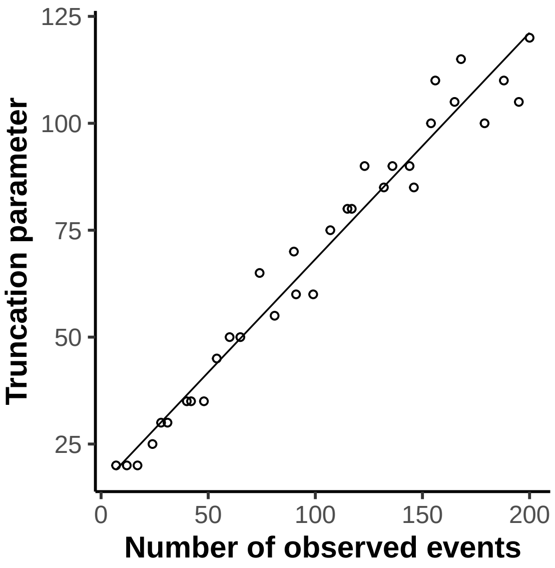


**Figure S2:** The truncation parameter required by the ODE approximation [MGVS20] grows approximately linearly with the size of the dataset. Each point in the scatter plot shows the size of the truncation parameter for a simulated dataset. The solid line shows a linear least squares fit.

#### Model validation and computational complexity

Our TimTam approximation was implemented in Haskell and the `criterion` library was used to estimate the average evaluation time (estimated by evaluating the log-likelihood for 5 seconds and counting the number of evaluations). For the ODE approximation we used the Cython implementation from [MGVS20] and used the Python Standard Library `timeit` module to estimate the average evaluation time (averaged over 50 replicates). We are most interested in the computational complexity of the algorithms in terms of the size of the dataset they are applied to. Because the implementations are in different languages, an absolute comparison is difficult (although since Cython and Haskell are used, they should give a reasonable indication of the performance that could be achieved with a low-level language).

To model the computational complexity of the likelihoods we fit a linear model to the logarithms of the evaluation times and the size of the dataset. Thus, if $n$ is the size of the dataset, the time to evaluate the log-likelihood, $t_{\text{eval}}$, we have $t_{\text{eval}} \propto n^a$. An estimate of $a \approx 1$ suggests a linear complexity and an estimate $\approx 2$ a quadratic complexity. The average evaluation times and the model fit are shown in Figure 3 of the main text.

Using TimTam, the estimated value of the exponent of the fitted model is 1.03 with a 95% confidence interval of $(1.02, 1.04)$. Using the ODE approximation it is 2.38 with a 95% confidence interval of $(2.26, 2.50)$. For both algorithms smaller datasets appear as outliers (likely due to the computational overhead of the programs.) We repeated the estimation process with robust linear regression (using `rlm` from the `MASS` package in R), under this model the exponent was 1.02, $(1.02, 1.03)$ for the TimTam likelihood and 2.49, $(2.39, 2.58)$ for the ODE approximation.

#### Simulation with all data types

To evaluate statistical power to identify model paramaters we simulated a dataset using the parameters listed in Table 1 of the main text with additional scheduled sampling events. The first unsequenced scheduled sampling event occurred at time $t = 2$ and then was repeated every 1.5 time units, the first sequenced scheduled event occurred at time $t = 2.5$ and was repeated every 1.5 time units. For all scheduled events the sampling probability was set to 0.05.

To show the effect of the simulation length and the number of observations on statistical power, we truncated the simulation at two timepoints, $t = 12$ and $t = 16$. Figure S3 shows cross-sections of the TimTam log-likelihood function generated by fixing the parameters to (a) their values from the simulation or (b) their MLE obtained numerically, while fixing $\mu$ at its true

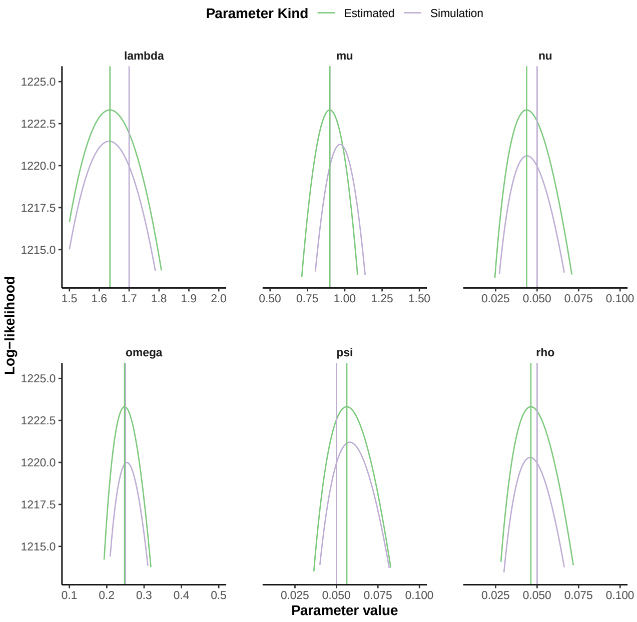


**Figure S3:** Cross sections of the log-likelihood function taken about the parameter values used in the simulation (lilac) and the estimated values (green), both of which are indicated by vertical lines, given the data that was available at $t = 16$.

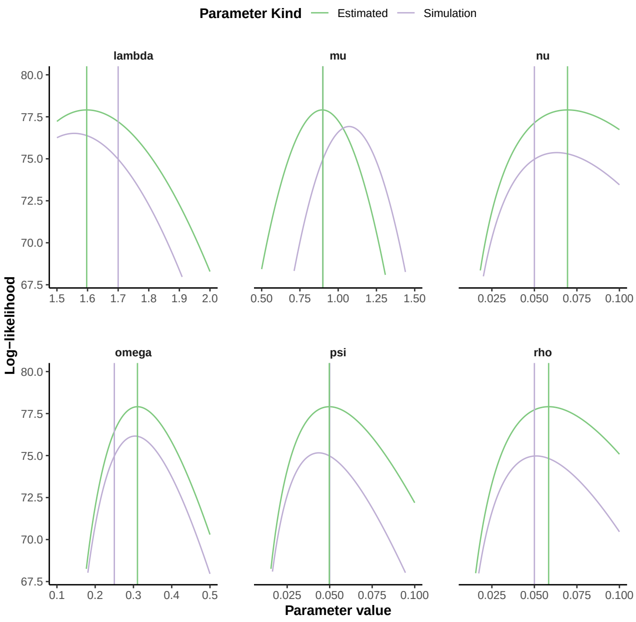


**Figure S4:** Cross sections of the log-likelihood function taken about the parameter values used in the simulation and the estimated values (both of which are indicated by vertical lines) given the data that was available at time 12.

value, and then varying each element of the parameter vector individually to explore the surface. The log-likelihood cross-sections for the datasets truncated at $t = 12$ is shown in Supplementary Figures S4.

We also investigated how well TimTam estimates the prevalence of hidden lineages through time. Figure S5 shows the prevalence in the simulation through time, together with the estimates of the prevalence made using the incomplete data available at that point in the simulation.

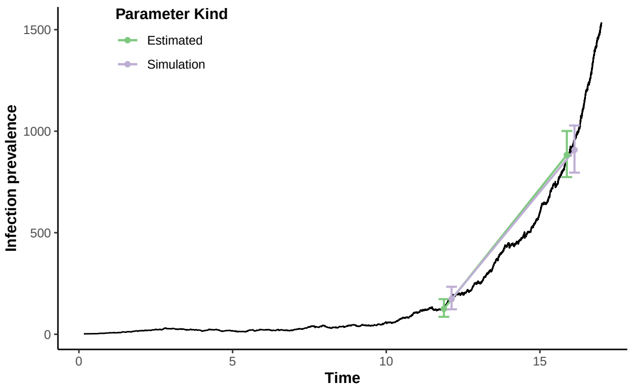


**Figure S5:** The LTT plot of the simulated dataset and the estimates of the prevalence generated at a couple points in the simulation (based on only the data prior to that point in time) using either the "true" parameters responsible for the simulated data or the estimated values conditional upon the data up to that point in the simulation.

#### Parameter identifiability and aggregation scheme

In the simulation study of the effects of aggregation of observations described in the main text we used MCMC to characterise the posterior distribution. For each dataset a single chain was run for $5 \times 10^4$ iterations using a Gaussian kernel with a standard deviation of 0.01. The first thousand samples were removed as burn-in and convergence was assessed via visual inspection of the traces. The effective sample size was greater than 200 and the standard error was substantially less than the posterior standard deviation for all parameters. Diagnostics were computed using `coda` [PBCV06].

The joint distribution of the posterior samples conditional upon the unscheduled dataset are shown in Figure S6 and for the scheduled dataset obtained via aggregation in Figure S7.

#### Repeated simulation to test credible interval coverage

A simulation study was carried out to test whether the 95% credible intervals (CI) of the birth rate, $\lambda$, and the prevalence at the present, $H(t_N)$, contain the values from the simulation. Since these are *credible intervals* (not confidence intervals) in general they will not contain the simulation parameters with the correct frequency. However, since we have used a uniform prior over the full support of the parameters, the posterior is just the likelihood. Consequently, in a large-sample setting, we would expect the CI to behave similarly to a confidence interval and contain the simulation parameters with approximately the correct frequency. The birth rate is used instead of the reproduction number because it is ill-defined in the case of the aggregated data. This is because there the scheduled events arise from post processing of the data, i.e., the binning of samples, instead of being generated by the sampling

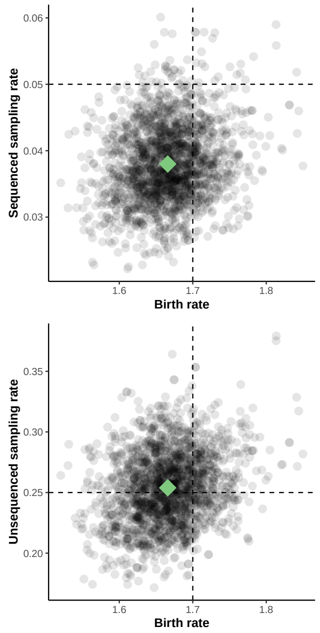


**Figure S6:** Given the death rate, $\mu$, the posterior distribution given scheduled observations has a well-defined maximum. The scatter plots show the joint posterior distribution of a subset of $10^3$ samples from the MCMC. The green point shows the posterior mean of the samples and the vertical and horizontal dashed lines show the true values used in the simulation of the dataset.

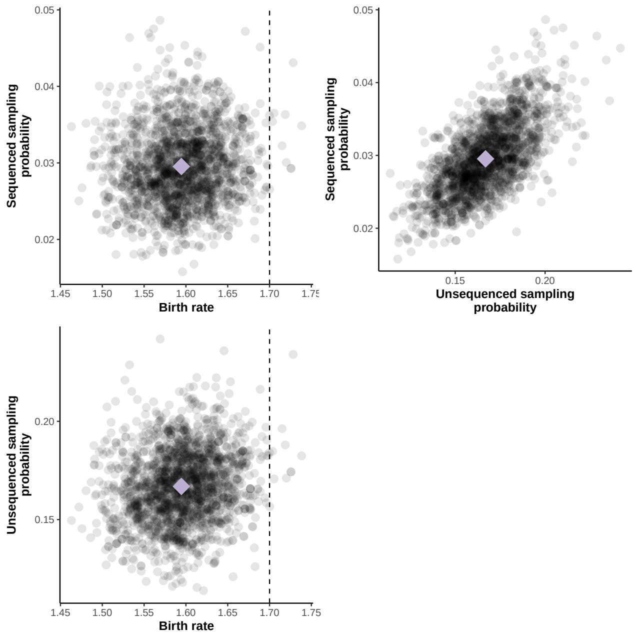


**Figure S7:** Given the death rate, $\mu$, the posterior distribution (from aggregated unscheduled observations) has a well-defined maximum. The scatter plots show the joint posterior distribution of a subset of $10^3$ samples from the MCMC. The lilac point shows the posterior mean of the parameter and the vertical lines show the true birth rate used in the simulation (there is no obvious equivalent for the other parameters).

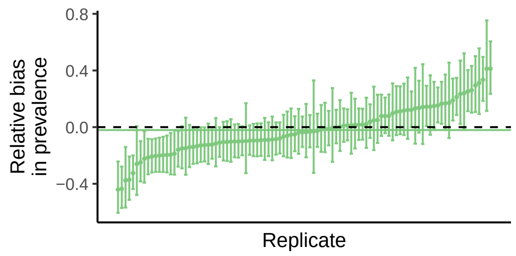


**(a)** Using simulated unscheduled events as data

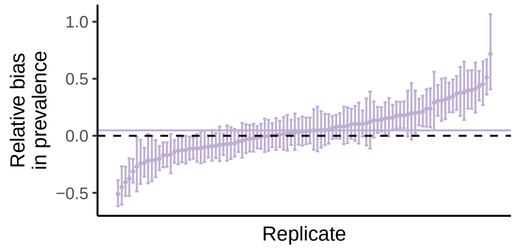


**(b)** Treating aggregated data as scheduled observations

**Figure S8:** The 95% range of relative bias in the estimates of the prevalence across the replicates. **(a)** The results using the simulated unscheduled observations. **(b)** The results when these unscheduled events are aggregated and the resulting aggregated data is treated as scheduled observations. The solid horizontal lines indicate the mean of the point estimates and the dashed line corresponds to zero bias.

model.

Figure S8 shows 95% CI for the relative bias in the estimate of the prevalence at the present. Of 100 replicates, 64 and 54 of the CIs contained the true prevalence from the simulation using the unscheduled and aggregated data respectively. Figure S9 shows 95% credible intervals for the estimate of the birth rate. Of the 100 replicates, 90 and 54 of the CIs contained the true birth rate from the simulation using the unscheduled and aggregated data respectively.

#### Bias due to aggregation

In Figure 5 of the main text we can see that the birth rate is underestimated when conditioning upon the aggregated data. Moreover, from Figure S9 we can see that this is not particular to that realisation of the process. That the bias is towards underestimation rather than overestimation is a consequence of the way in which the observations have been aggregated. Had they been aggregated into a single event at the start of the interval, then the bias would be reversed. To see why this is the case, consider an interval of time $[0, T]$, where a unscheduled, unsequenced observation (an $\omega$-sample) occurs at time $T/2$. Under the existing aggregation scheme, this sample will be treated as if it was removed from the population at time $T$ even though it had been removed at time $T/2$. This means that there is $T/2$ units of time where it has been removed from the population but the likelihood treats it as existing in an infectious state. Adjusting for the infections the removed individual did not generate is why lower values of the birth rate, $\lambda$, have an increased likelihood.

If aggregating at the start of the interval leads to overestimation, and aggregating at the end of the interval underestimation, one might consider aggregating

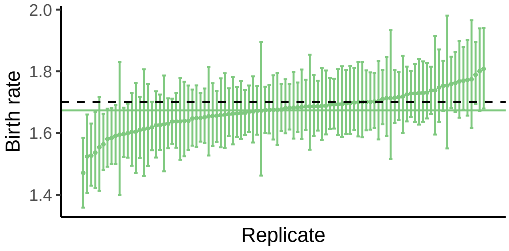


**(a)** Using simulated unscheduled events as data

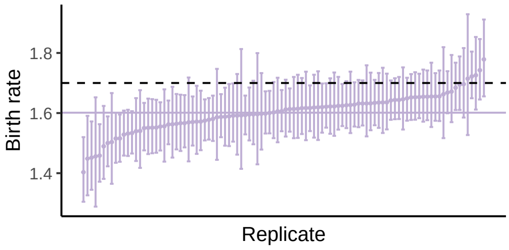


**(b)** Treating aggregated data as scheduled observations

**Figure S9:** The 95% credible intervals for the birth, $\lambda$, across the replicates. **(a)** The results using the simulated unscheduled observations. **(b)** The results when these unscheduled events are aggregated and the resulting aggregated data is treated as scheduled observations. The solid horizontal lines indicate the mean of the point estimates and the dashed line the true value of 1.7.

at an intermediate time. Considering the mean field approximation for when unscheduled events occur one can derive a midpoint:

$$t_{\text{agg}} = \frac{1}{\lambda - \mu - \omega - \psi} \log\left(\frac{1}{2}\left[e^{(\lambda-\mu-\omega-\psi)T} + 1\right]\right)$$

as the time when half the unscheduled events will be shifted forwards in time during the aggregation and half will be shifted backwards.

### Literature Cited

[FS09] Philippe Flajolet and Robert Sedgewick. *Analytic Combinatorics*. Cambridge University Press, 2009.

[MGVS20] Marc Manceau, Ankit Gupta, Timothy Vaughan, and Tanja Stadler. The probability distribution of the ancestral population size conditioned on the reconstructed phylogenetic tree with occurrence data. *Journal of Theoretical Biology*, page 110400, 2020.

[PBCV06] Martyn Plummer, Nicky Best, Kate Cowles, and Karen Vines. CODA: Convergence Diagnosis and Output Analysis for MCMC. *R News*, 6(1):7–11, 2006.

[Sta10] Tanja Stadler. Sampling-through-time in birth-death trees. *Journal of Theoretical Biology*, 267(3):396–404, 2010.
